# PAGE4 and Conformational Switching: Insights from Molecular Dynamics Simulations and Implications for Prostate Cancer

**DOI:** 10.1101/264010

**Authors:** Xingcheng Lin, Susmita Roy, Mohit Kumar Jolly, Federico Bocci, Nicholas Schafer, Min-Yeh Tsai, Yihong Chen, Yanan He, Alexander Grishaev, Keith Weninger, John Orban, Prakash Kulkarni, Govindan Rangarajan, Herbert Levine, José N. Onuchic

## Abstract

Prostate-Associated Gene 4 (PAGE4) is a disordered protein implicated in the progression of prostate cancer. PAGE4 can be phosphorylated at two residue sites by Homeodomain-Interacting Protein Kinase 1 (HIPK1) to facilitate its binding to the Activator Protein-1 (AP-1) transcription factor. In contrast, a further hyperphosphorylation of PAGE4 by CDC-Like Kinase 2 (CLK2) reduces its binding affinity to AP-1, thus affecting the androgen receptor (AR) activity. Both SAXS and smFRET experiments have shown a structural expansion of PAGE4 upon hyperphosphorylation and a significant increase in size at its N-terminal half than that at its C-terminus. To understand the molecular mechanism underlying this structural transition, we performed a series of constant temperature molecular dynamics simulations using Atomistic AWSEM — a multi-scale molecular model combining detailed atomistic and coarse-grained simulation approaches. Our simulations show that electrostatic interaction drives a transient formation of an N-terminal loop, which causes the change in size for different phosphorylated forms of PAGE4. Phosphorylation also changes the preference of secondary structure formation of PAGE4, which signifies a transition between states that display different degree of disorder. Finally, we construct a mechanism-based mathematical model that allows us to capture the interactions of different forms of PAGE4 with AP-1 and AR, a key therapeutic target in prostate cancer. Our model predicts intracellular oscillatory dynamics of HIPK1-PAGE4, CLK2-PAGE4 and AR activity, indicating phenotypic heterogeneity in an isogenic cell population. Thus, conformational switching among different forms of PAGE4 may potentially affect the efficiency of therapeutic targeting of AR.

## 1 Introduction

It is now evident that a large fraction of the human proteome comprises of proteins, or regions within proteins, that lack stable structure under physiological conditions [1, 2, 3, 4]. Such unstructured proteins exist in large ensemble of interconverting conformations in order to function and thus, they are referred to as intrinsically disordered proteins (IDPs). Despite lack of rigid structure, IDPs are involved in regulation, signaling, and control of information flow in the system, where binding to multiple partners and high-specificity/low-affinity interactions play a crucial role [5, 6]. Furthermore, disorder is a unique structural feature that enables IDPs to participate in both one-to-many and many-to-one signaling [3, 7]. But when overexpressed, or aberrantly expressed, IDPs are prone to engage in promiscuous interactions and lead to different pathological states [8, 9].

IDPs in general have remained intractable to classical experimental methods and therefore, mechanistic insight into how they function is limited to a few IDPs at best. PAGE4 is an IDP that has been well characterized both structurally and functionally. It has the hallmarks of a proto-oncogene; while it is highly expressed during its development in the fetal prostate [10, 11], it is aberrantly expressed in the diseased gland where it plays an important role in tumorigenesis [10].

PAGE4 is a stress-response factor [10]; it functions as transcriptional coactivator and potentiates transactivation by c-Jun [11], a component of the AP-1 transcription factor complex that can negatively regulate the activity of the androgen receptor (AR) in prostate cancer (PCa) cells [12, 13]. Furthermore, in PCa cells, PAGE4 is phosphorylated by the stress-response kinase Homeodomain-Interacting Protein Kinase 1 (HIPK1) predominantly at Thr51 and to a significantly lower level at Ser9 [14] but is hyperphosphorylated by a second kinase, namely CDC-Like Kinase 2 (CLK2) [15]. Cell-based studies reveal that phosphorylation of PAGE4 by the two kinases leads to opposing functions. HIPK1-phosphorylated PAGE4 (HIPK1-PAGE4) potentiates c-Jun but CLK2-phosphorylated PAGE4 (CLK2-PAGE4) attenuates c-Jun activity [15]. Biophysical measurements employing small-angle X-ray scattering (SAXS), single-molecule fluorescence resonance energy transfer (smFRET), and NMR indicate that HIPK1-PAGE4 exhibits a relatively compact conformational ensemble that binds AP-1, whereas CLK2-PAGE4 is more expanded and resembles a random coil with reduced affinity for AP-1 [15]. Although the data indicated different sizes and functions of PAGE4 with increasing phosphorylation, the atomistic details of critical interactions that are responsible for such conformational changes remain unclear.

To investigate the structural details causing the change of size of PAGE4 upon phosphorylation, and to overcome the time resolution limit associated with these biophysical experiments, molecular dynamics (MD) simulations with a physical-based model are needed as a complementary effort. One molecular dynamics approach based on the funneled energy landscape theory [16, 17, 18] is the associative memory, water mediated, structure and energy model (AWSEM) [19]. AWSEM has been effective not only in the correct prediction of protein structures and binding interfaces [19, 20], but also in elucidating the molecular mechanisms underlying protein folding and regulation of cellular responses [21]. An important component of the AWSEM force field, called an associated memory term, utilizes local structural signals from external sources, such as protein fragments sharing similar sequence information in the protein database [22].

Recently, we developed a multi-scale atomistic AWSEM model (AAWSEM) [23], which harnesses the power of all-atom explicit-solvent simulations to directly generate atomistic structures for use as associative memory terms of an AWSEM simulation. AAWSEM has been shown to be able to reliably fold both *α*-exclusive proteins [23] and *α/β* proteins [24]. In particular, AAWSEM is especially useful for investigating dynamics of disordered proteins such as PAGE4 because the lack of known structure limits the utility of the AWSEM model. In addition, the coarse-grained nature of AAWSEM is suitable for fully sampling the conformational space of PAGE4 dynamics, which is significantly larger for disordered proteins compared to proteins in ordered states.

Here, we performed simulations with AAWSEM to investigate the mechanism(s) underlying structural transitions of the PAGE4 conformational ensemble. Consistent with the experimental findings, our simulations produce a change of size in PAGE4 upon phosphorylation. In addition, we suggest that formation of an N-terminal loop is the source of this variation. After gaining insights into conformational dynamics of different forms of PAGE4 (Wild type un-phosphorylated WT-PAGE4, double-phosphorylated HIPK1-PAGE4 and hyper-phosphorylated CLK2-PAGE4), we developed a mechanism-based mathematical model to describe how switching from one form to another (e.g. from HIPK1-PAGE4 to CLK2-PAGE4) can affect the dynamics of the PAGE4/AP-1/AR regulatory circuit. The oscillations observed in this circuit suggest that PAGE4 conformational dynamics may contribute to phenotypic plasticity, and thus potentially the efficacy of therapeutic targeting of AR in PCa.

## 2 Results

### 2.1 The sequence of PAGE4

PAGE4 is a 102-residue disordered protein. Most of the residues are predicted to have > %50 chance to be a random coil by secondary structure prediction tools [25] (except for residue Cys63 and Gln64, which are predicted to be helical and located in the region experimentally found to go transiently into helix) (Figure 1). PAGE4 is also a highly charged protein, with a total of 16 positively charged residues and 24 negatively charged residues, resulting in a net charge of −8. The charged residues are not evenly distributed. The N- (residue ID 4 to 12) and C- (residue ID 82 to 95) motifs are positively charged, and there is a central acidic region (residue ID 43 to 62) that is negatively charged. Also, there is a nine-residue region (residue ID 65 to 73) which can transiently turn into helix as shown by NMR experiment [26]. HIPK1 phosphorylates two sites in PAGE4, with a partial phosphorylation at Ser9 (30% ∼ 40% chance, within the N-motif) and a complete phosphorylation of Thr51 (> 95% chance, within the central acidic region) [26]. CLK2 phosphorylates 8 sites of PAGE4, with most of them completely phosphorylated: Ser7 (> 95%), Ser9 (> 95%), Thr51 (> 95%), Thr71 ( 50%), Ser73 ( 75%), Ser79 ( 50%), Thr85 (> 95%) and Thr94 ( 60%) [15]. Since both Ser7 and Ser9 reside in the N-motif, and Thr85 and Thr94 reside in the C-motif, the hyperphosphorylation neutralizes the N- and C motifs. On the other hand, Thr51 resides in the central acidic region, and Thr71 and Ser73 reside in the transient helix region. The phosphorylation of these amino acids will likely change the local arrangement of residues around this region, contributing to the shift of structural ensemble.

**Figure 1:**
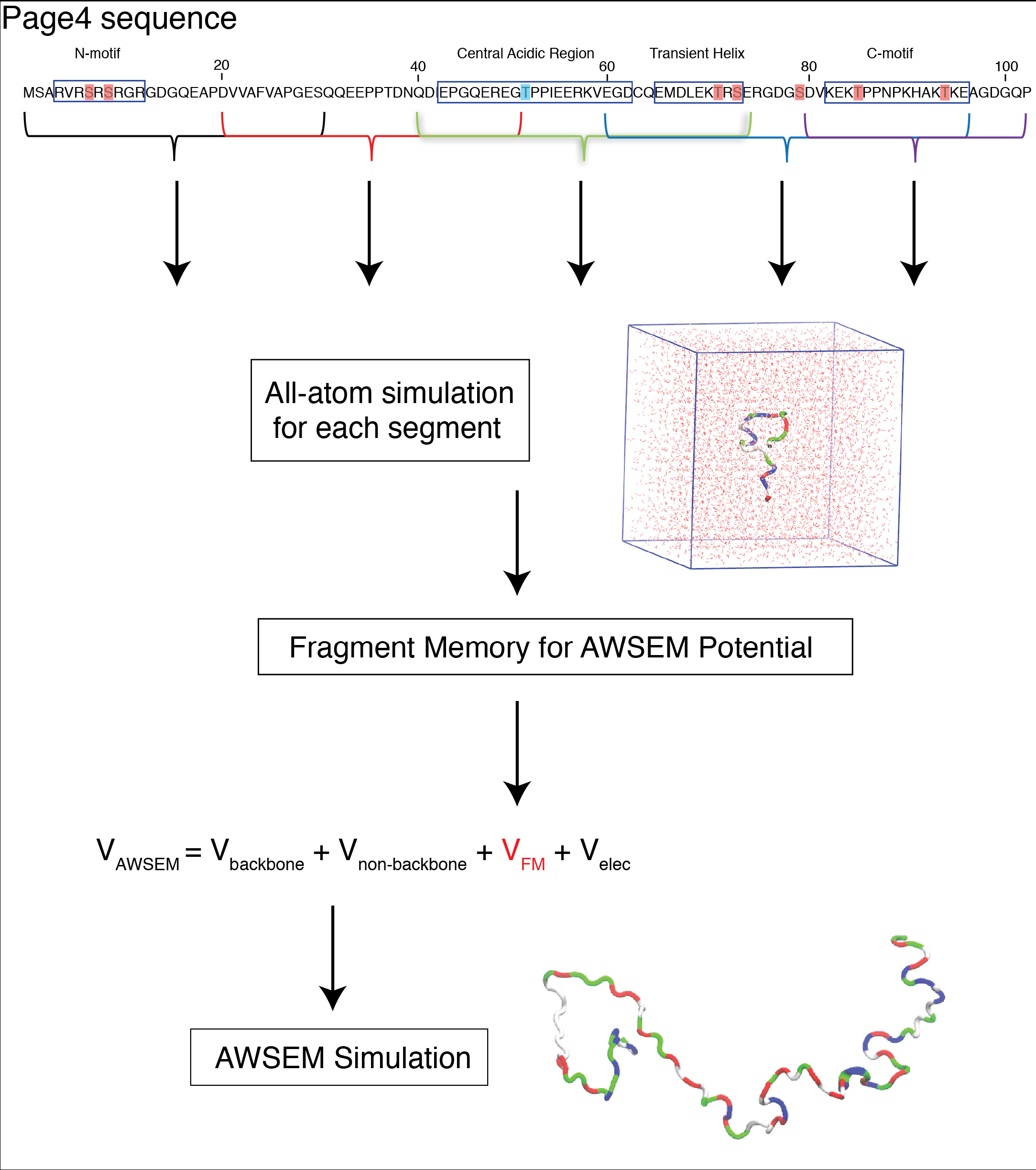
The sequence nature of PAGE4 and the flowchart for the AAWSEM simulation. The PAGE4 sequence was adapted from [15]. The phosphorylation sites in HIPK1-PAGE4 are highlighted in blue, and those in CLK2-PAGE4 are highlighted in red. Each phosphorylated form of PAGE4 protein was separated into 5 segments according to its sequence nature: Segment 1 (residue ID 1 to 30), Segment 2 (residue ID 20 to 50), Segment 3 (residue ID 40 to 74), Segment 4 (residue ID 60 to 96) and Segment 5 (residue ID 80 to 102). Each segment was simulated in a explicit-solvent environment with Continuous Simulated Tempering (CST) [27]. The structural ensemble from CST at T = 293 ∼ 330K was clustered into structural ensemble and passed as fragment memories into AWSEM model for the coarse-grained level simulations of the entire PAGE4 (a snapshot of which in simulation is colored by residue type in the right bottom).

### 2.2 The Phosphorylation by HIPK1 and CLK2 Changes the Size of PAGE4

Our simulations indicate distinct changes in the size of PAGE4 upon different levels of phosphorylation. The size of the protein can be measured by the radius of gyration (*R_g_*) of the protein during these simulations. The preference of the size of the protein structures in our simulation can be captured by a free energy plot (Figure 2 A). The free energy *F* was calculated as *F* = −*k_B_Tlog(P)* where *P* is the probability for the proteins to be at a specific *R_g_*. Thus, the lowest free energy in this plot corresponds to the *R_g_* of the most populated ensemble in the conformational space. It is clear that CLK2-PAGE4 is more expanded compared with HIPK1-PAGE4, which is also slightly more compact than WT-PAGE4.

**Figure 2:**
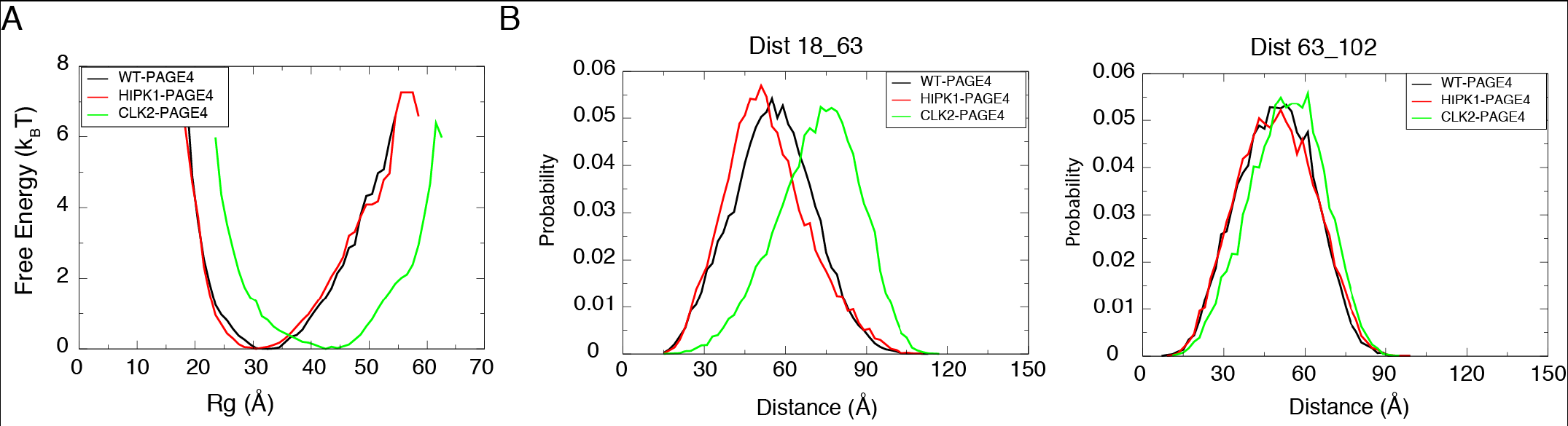
**A**. The free energy plot based on the Radius of Gyration (*R_g_*) of the simulated proteins. **B**. The probability distribution of distance between two residue pairs that were previously measured in smFRET experiments.

In order to quantify the size of different forms of PAGE4 — WT-PAGE4, HIPK1-PAGE4 and CLK2-PAGE4 — and to compare them with experimental measurements [15], we calculated the average *R_g_* of each form of PAGE4 following equation (1), as given below:

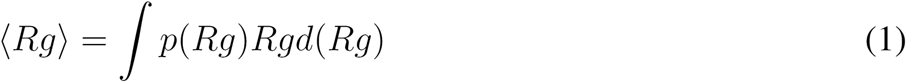

where *p(R_g_)* is the probability for the protein to be at a specific *R_g_* value. The calculated *R_gs_* for three forms of PAGE4 are presented in the second column of Table 1. The simulations qualitatively reproduce the change of size upon phosphorylation, as seen in the SAXS experiments [15]. Moreover, the calculated 〈*R_g_*〉 is also quantitatively similar to the experimental values (Table 1).

**Table 1:** A summary of the results from the simulations of different forms of PAGE4.

| | $\langle Rg(EXP) \rangle$<br>$^a$ (Å) | $\langle Rg(SIM) \rangle$<br>$^b$ (Å) | RMS Dist<br>18_63<br>(EXP) (Å) | RMS Dist<br>18_63<br>(SIM) (Å) | RMS Dist<br>63_102<br>(EXP) (Å) | RMS Dist<br>63_102<br>(SIM) (Å) |
| --- | --- | --- | --- | --- | --- | --- |
| Wildtype<br>Form | $36 \pm 1.1$ | 32.9 | 56 | 57.4 | 50 | 51.2 |
| HIPK1<br>Form | $34.7 \pm 1.2$ | 32.1 | 59 | 55.6 | 50 | 51.4 |
| CLK2<br>Form | $49.8 \pm 1.9$ | 41.8 | 75 | 73.4 | 55 | 54.8 |

<sup>a</sup>EXP stands for results from experiments [15]
<sup>b</sup>SIM stands for results from simulations

### 2.3 Phosphorylation expands the structure at the N-terminal end of PAGE4 more than that of the C-terminus

Aside from the SAXS experiments, the smFRET experiments helped us gain further insight into the expansion of PAGE4 upon phosphorylation [15]. The experiment found that the size of the N-terminal half of the protein, manifested as the distance between the donor and acceptor dyes tagged at the residues 18 and 63 (Dist18_63), had a more significant increase of size in the hyper-phosphorylated form (56 to 75 Å) than in its double-phosphorylated variant (56 to 59 Å). In comparison, the size of the C-terminal PAGE4 is less sensitive to phosphorylation. We find that the distance between residue 63 and 102 (Dist63_102) changes from 50 to 55 Å (Table 1) upon hyperphosphorylation, while we do not observe any noticeable change upon double-phosphorylation.

Our simulations reproduce distances that are similar to experimental observations (Table 1). Since our coarse-grained simulation model uses three beads (*C_α_*, *C_β_* and O atoms) to represent a residue, we approximated the distance between the probes of two residues as the distance between their *C_α_* atoms. Consistent with the observations, our simulations show that Dist18_63 changes from 57.4 Å in wildtype to 73.4 Å in CLK2-PAGE4, while Dist63_102 changes from 51.4 Å to 54.8 Å.

### 2.4 The structural expansion in the N-terminus is associated with the loss of a loop formation

MD simulations enable us to probe the structural details of PAGE4 dynamics. The collected ensembles of simulated structures show that the expansion of PAGE4 upon hyperphosphorylation is caused by the loss of a N-terminal loop formation between the N-Motif and the central acidic region (Figure 3). Structurally, the N-motif has an excess of basic residues in both WT-PAGE4 and HIPK1-PAGE4 to allow the N-motif to easily interact with the central acidic region around residue 40. These specific interactions lead to frequent loop formation. In contrast, the N-motif is neutralized upon hyperphosphorylation, which causes a loss of this loop formation and ultimately, a more expanded CLK2-PAGE4. To visualize this loop formation, we plot the average contact maps of proteins during their simulations (Figure 3). Because the electrostatic interactions were simulated in a Debye-Huckel form with a Debye length of 10.0 Å (see Method Section 4.1 for more details), we used the cutoff value of 20.0 Å between the *C_β_* atoms for creating this contact map. This choice of cut-off value captures the electrostatic interactions between residues before they decay to 20% of its value at Debye length. Also, since we are mostly interested in non-local interactions, contacts are defined only for the residues that are at least three residues apart from each other in sequence. As shown in Figure 3B, in both WT-PAGE4 and HIPK1-PAGE4, the N-motif region has a non-zero probability to form contacts with the central acidic region, while no contacts are formed in CLK2- PAGE4. There are also interactions being formed between the C-motif and the central acidic region in WT-PAGE4 and HIPK1-PAGE4, though the probability is smaller.

**Figure 3:**
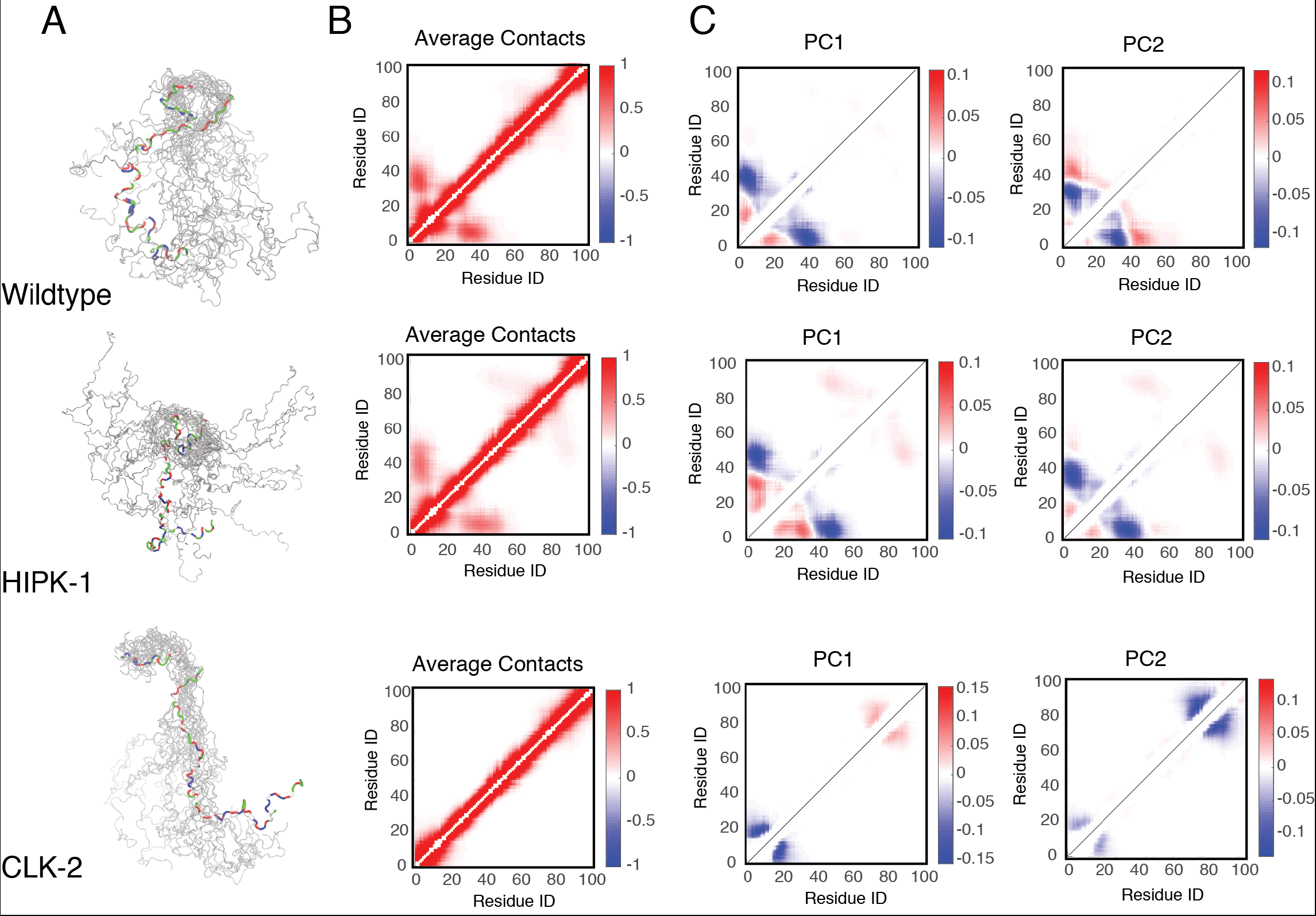
The structure ensemble of different forms of PAGE4 collected from the simulations. **A.** The structural ensemble generated during our simulations. Randomly picked structures are RMSD aligned by the N-motif [28]. It is clear that the N-terminal loop forms in the wildtype and HIPK1 form, while no loops are formed in the CLK2 form. **B**. The average contact maps generated based on the simulations. The color bar shows the probability of contact formation during our simulations. There are non-zero probabilities for interactions between the N-, C-motif and the central acidic region. **C**. The top three principal modes generated from the contact-based principal component analysis. We plot the coefficients of the first two principal components, which correspond to the collective change of each contact.

Although the averaged contact map demonstrates a higher likelihood for the N-terminal loop that forms in WT-PAGE4 and HIPK1-PAGE4 than in CLK2-PAGE4, it lacks a direct description of any dynamic features. To understand more regarding collective conformational movements, we used a Principal Component Analysis (PCA) calculated based on the contacts to probe the dominant mode of dynamics, motivated by the use of similar approaches before [29, 30]. We plot the coefficients of the top two principal components, which correspond to the variation of each contact within the simulated ensembles. A larger value of coefficient means more variation of the corresponding contact movement in that principal mode. In WT-PAGE4 and HIPK1-PAGE4, the calculated PC1 shows a collective motion of the N-motif towards (or away from) the central acidic region (the blue halo between residues ∼ 10 and residues ∼ 50 in Figure 3C), indicating a frequent loop formation in the N-terminal end of the proteins. In contrast, in CLK2-PAGE4, regular motions only happen among residues local in sequence. Interestingly, we also observe collective motion between the C-motif and the central acidic region in HIPK1-PAGE4. The principal component coefficients of this motion have the opposite sign to that of the N-motif (the red halo between residues ∼ 50 and residues ∼ 90 in Figure 3C), albeit much weaker than that of the N-motif. The observed PCA patterns thus suggest some competitive motions of the N, C-motif, which take turns to form a loop with the central acidic region. The observation of loop formation also hints at a fly-casting motion frequently observed in the study of disordered proteins [31, 32, 33].

### 2.5 Simulations find changes in turn-like structures upon phosphorylation

In addition to indicating the expansion of protein size upon hyperphosphorylation, NMR experiments suggested that the more compact structure of HIPK1-PAGE4 is accompanied by an increase of turn-like structures in the central acidic region [26]. Our simulations indeed find an increase of turn structure in this region also (Figure 4). The most prominent increase in turn structure occurs between Arg48 and Thr51, where the probability of forming a turn structure increases from 40% to 80% (Figure 4A). This change is probably due to the phosphorylation of Thr51. The simulations also display a more populated turn structure in the same region upon hyperphosphorylation. Because no other residues close to Thr51 are further phosphorylated upon hyperphosphorylation, CLK2-PAGE4 shares the same structural features as HIPK1-PAGE4 in the central acidic region.

**Figure 4:**
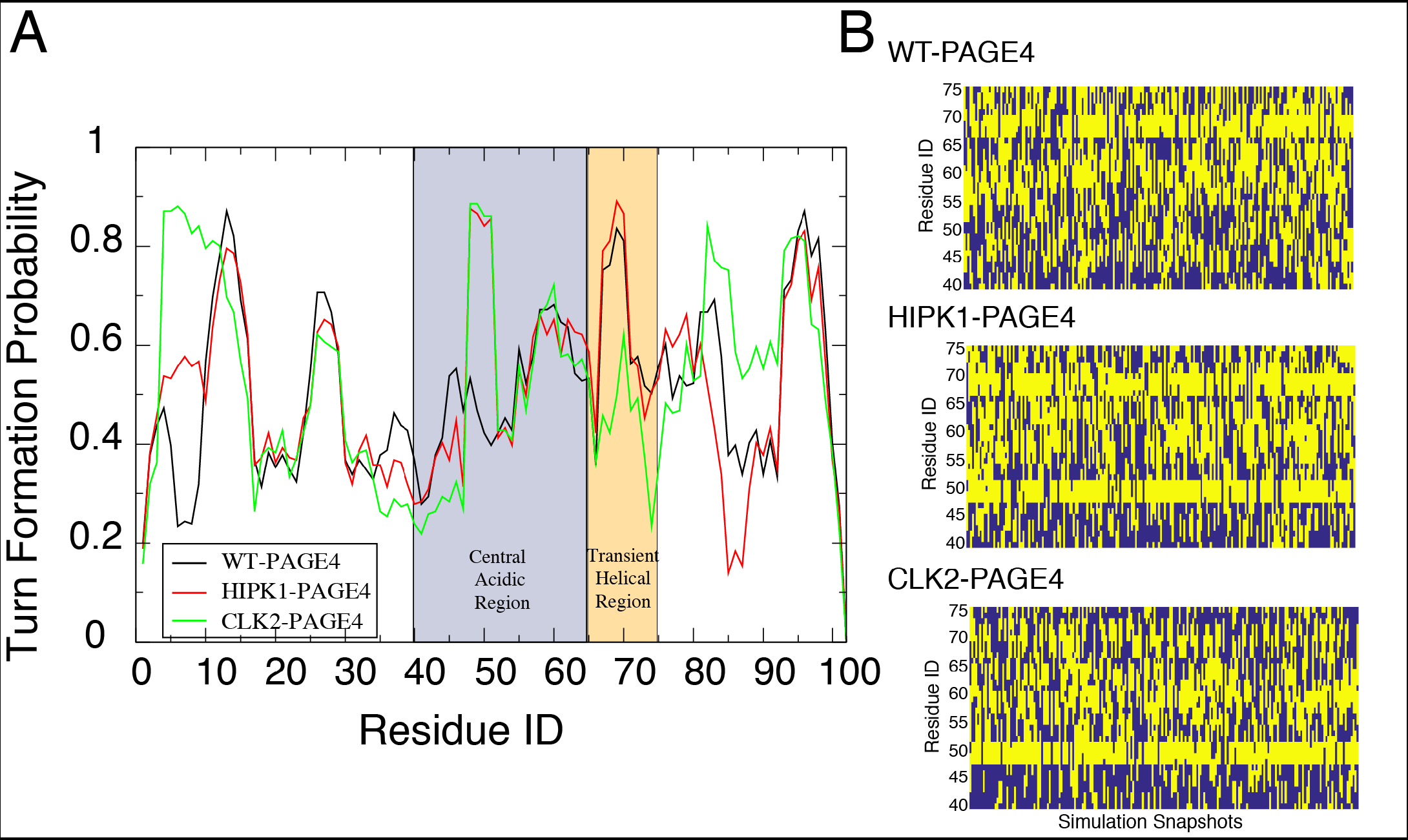
**A**. The probability for each residue to be a turn-like structure. The central acidic region and transient helical region are shaded in blue and orange respectively. The secondary structure was calculated using Stride algorithm [34]. **B**. The dynamics of turn formation per residue along the simulations of each phosphorylated form of PAGE4. We only show the residues in the central acidic region and the transient helical region. Yellow means the residue is identified by Stride algorithm to be a turn structure, while blue means it is not.

Two other residues in the transient helix region (Thr71 and Ser73) are further phosphorylated by CLK2. Phosphorylation of these residues causes a reduction of turn structure in this region (Figure 4A). The simulation time trace of turn-like structure in this region also changes upon phosphorylation (Figure 4B). The change of turn-like structure signifies a transient order-disorder transition in this local region. It is hypothesized that the binding site for the AP-1 complex is located in this region of PAGE4 [26]. Therefore, our simulations suggest that the hyperphosphorylation of PAGE4 by CLK2 results in a loss of order in the transient helix, which may then be responsible for a decrease in binding affinity between PAGE4 and the AP-1 complex.

### 2.6 Implications of switching between CLK2-PAGE4 and HIPK1-PAGE4 for plasticity in prostate cancer cells

To investigate the consequences of PAGE4 structural dynamics in modulating cellular plasticity, we constructed a coarse-grained mathematical model for PAGE4 intracellular dynamics and its implications in interacting with the AP-1/AR signaling axis (Figure 5A). The model considers the three relevant PAGE4 forms (WT-PAGE4, HIPK1-PAGE4 and CLK2-PAGE4) and the enzymes catalyzing the reactions among them (HIPK1, CLK2). As mentioned above, HIPK1-PAGE4 and CLK2-PAGE4 have significant differences in their potentiation of c-Jun and binding to AP-1. Because c-Jun potentiation can indirectly increase the levels of CLK2, a negative feedback loop comes into play that can give rise to oscillatory behavior for the levels of HIPK1-PAGE4, CLK2-PAGE4 and consequently AR activity [15]. This PAGE4/AP-1/AR circuit can generate oscillations between an androgen-dependent phenotype (characterized by high levels of HIPK1-PAGE4) and an androgen-independent phenotype (characterized by high levels of CLK2-PAGE4) [15]. Here, we extended this model by including the temporal dynamics of WT-PAGE4 and modeling different hormone-deprivation treatments for prostate cancer (see Methods section 4.5 for further details). The extended model exhibited similar phenotypic oscillations between a (high HIPK1-PAGE4, low CLK2-PAGE4) state and a (low HIPK1-PAGE4, high CLK2-PAGE4) state when no treatment was applied (Figure 5B) in quite a robust manner; a 10-fold variation of model parameters also exhibited oscillations (Figure S3 A-B).

**Figure 5:**
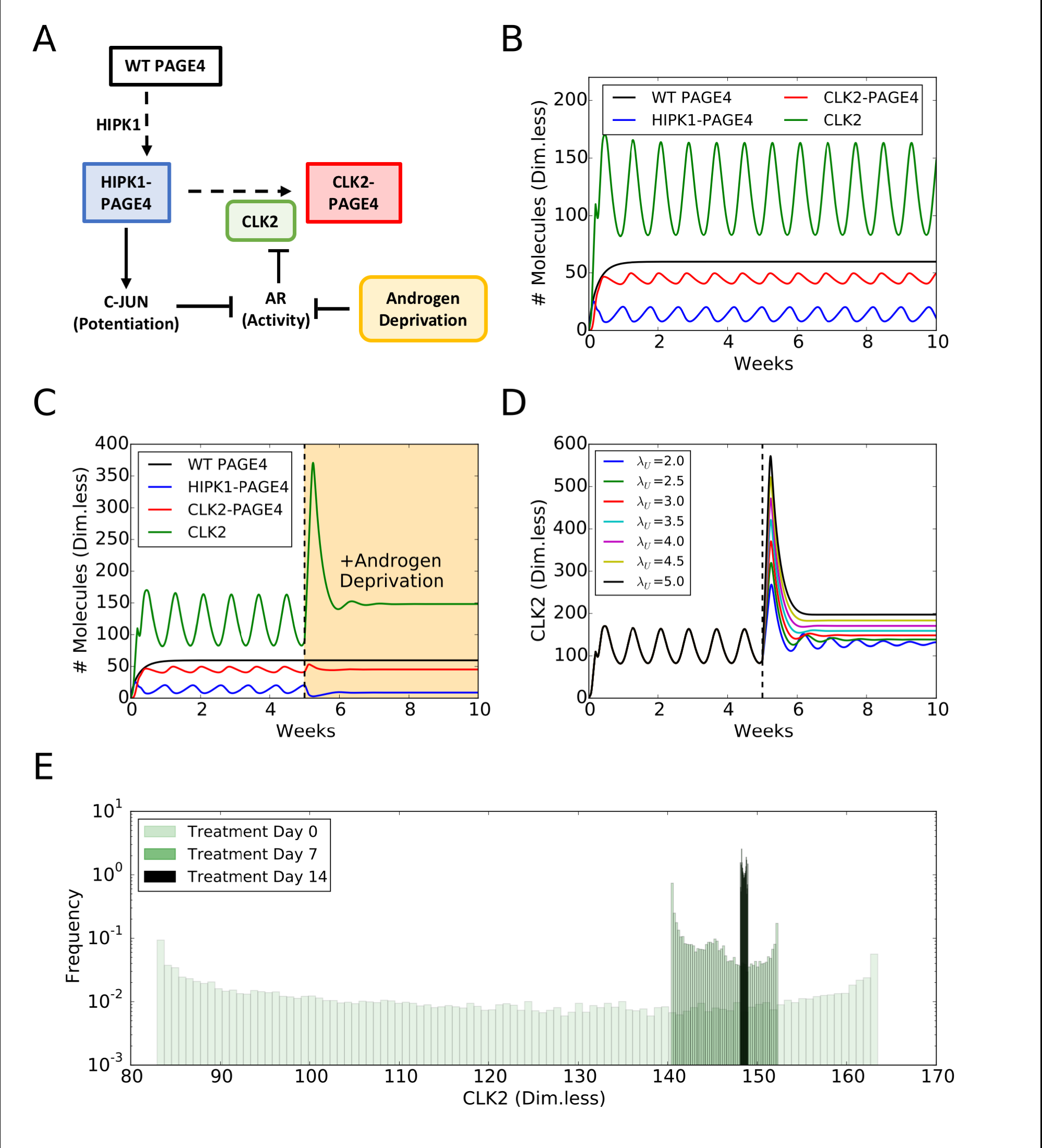
Androgen deprivation suppresses phenotypic oscillations and eliminates heterogeneity in a prostate cancer cell population, as predicted by a mathematical model. **A.** Schematic of PAGE4 phosphorylation circuit and its connection with androgen resistance. Wild-type PAGE4 is double phosphorylated by HIPK1 enzyme, and the HIPK1-PAGE4 complex is hyperphosphorylated via CLK2 enzyme. Further, HIPK1-PAGE4 regulates CLK2 via the intermediates C-Jun and Androgen Receptor (AR). The cell under hormonal therapy is deprived of androgen, thereby decreasing AR activity. **B.** Temporal dynamics of the cellular level of WT PAGE4, HIPK1-PAGE4, CLK2-PAGE4 and CLK2. Without ADT, the oscillatory behaviour exhibits a period of approximately 1 week. **C.** ADT (orange-shaded area) quenches oscillations within approximately 2 weeks. **D.** Temporal dynamics of the CLK2 cellular level for various CLK2 production rate fold-changes Au due to ADT. Any fold-change larger/equal than 2 is sufficient to quench the oscillations. The value used in (C) is 3. **E.** Distribution of CLK2 cellular level in a simulated cohort of 10000 prostate cancer cells. Before treatment (Day 0), the distribution spans over the whole range of CLK2 levels observed in the single cell simulation of (B). After one week of treatment (Day 7), the distribution considerably shrinks. At Day 14 of treatment, all cells have a similar level of CLK2. In all plots, WT PAGE4, HIPK1-PAGE4, CLK2-PAGE4 and CLK2 are represented in arbitrary dimensionless units.

First, we establish the period of oscillation. The relatively large half-life of HIPK1-PAGE4 (≈ 150 hours) [14] enables an oscillation period of approximately a week (Figure 5B). This timescale is comparable with various hormone-deprivation treatments applied in the clinic [35, 36] and motivated investigating the effect of various therapies. As a first step, we considered a constant inhibitory signal acting on the Androgen Receptor (AR), to simulate the effect of continuous Androgen Deprivation Therapy (ADT) — the standard care of therapy for over 75 years for locally advanced and/or metastatic PCa [35]. ADT quenched the oscillations and pushed the cell to a resistant phenotype (Figure 5C). This result depended on the assumed fold-change in the decrease of AR activity levels, but depended weakly on the specific mathematical form chosen to model ADT (Figure 5D, S3 C-D).

Next, we simulated continuous ADT in a cohort of 10000 independent prostate cancer cells. Each cell displays oscillations with the same period, but they are not synchronized, i.e. their oscillations have a non-zero phase difference. Therefore, the distribution of intracellular levels of CLK2 is quite broad before ADT is applied (Figure 5E, Day 0). After a week of therapy, the heterogeneity is considerably decreased, and after 2 weeks, all cells present similar levels (compare the distribution of CLK2 levels in Day 0, Day 7 and Day 14 in Figure 5E). Thus, ADT for a period of 2 weeks can not only quench the oscillations, but also limit phenotypic heterogeneity significantly in a prostate cancer cell population. It should be noted that our model focuses on cells that are not killed by therapeutic treatment. Thus, on one hand, cells with an androgen-dependent phenotype are highly likely to be killed by ADT, and on the other, ADT limits phenotypic heterogeneity among the cells that are not killed by ADT.

Motivated by these results, we implemented an Intermittent ADT [35] where cells are periodically exposed to ADT for a limited time (2 weeks on — 2 weeks off, in the simulation). Oscillations disappeared during the cycles of ADT, but emerged once ADT was removed (Figure 6A). Intriguingly, intermittent ADT coupled the oscillations of unsynchronized cells. We set up a simulation of 4 unsynchronized cells and tracked their level of CLK2-PAGE4. The oscillations became synchronized after the first round of ADT because the treatment pushed all cells to a similar state (Figure 6B). Further, we varied the duration of ADT pulses, and discovered that a single duration as quick as 4 days could synchronize cells in the model (Figure S4). Such synchronization can limit the phenotypic heterogeneity in cells that survive androgen deprivation. Intermittent Androgen Deprivation (IAD) therapy has been studied both in xenografts and in the clinic, with varying time-periods of androgen deprivation and the drug holiday in between [37]. Our model simulations offer a plausible explanation for why the period of androgen deprivation should be at least on the order of a few weeks.

**Figure 6:**
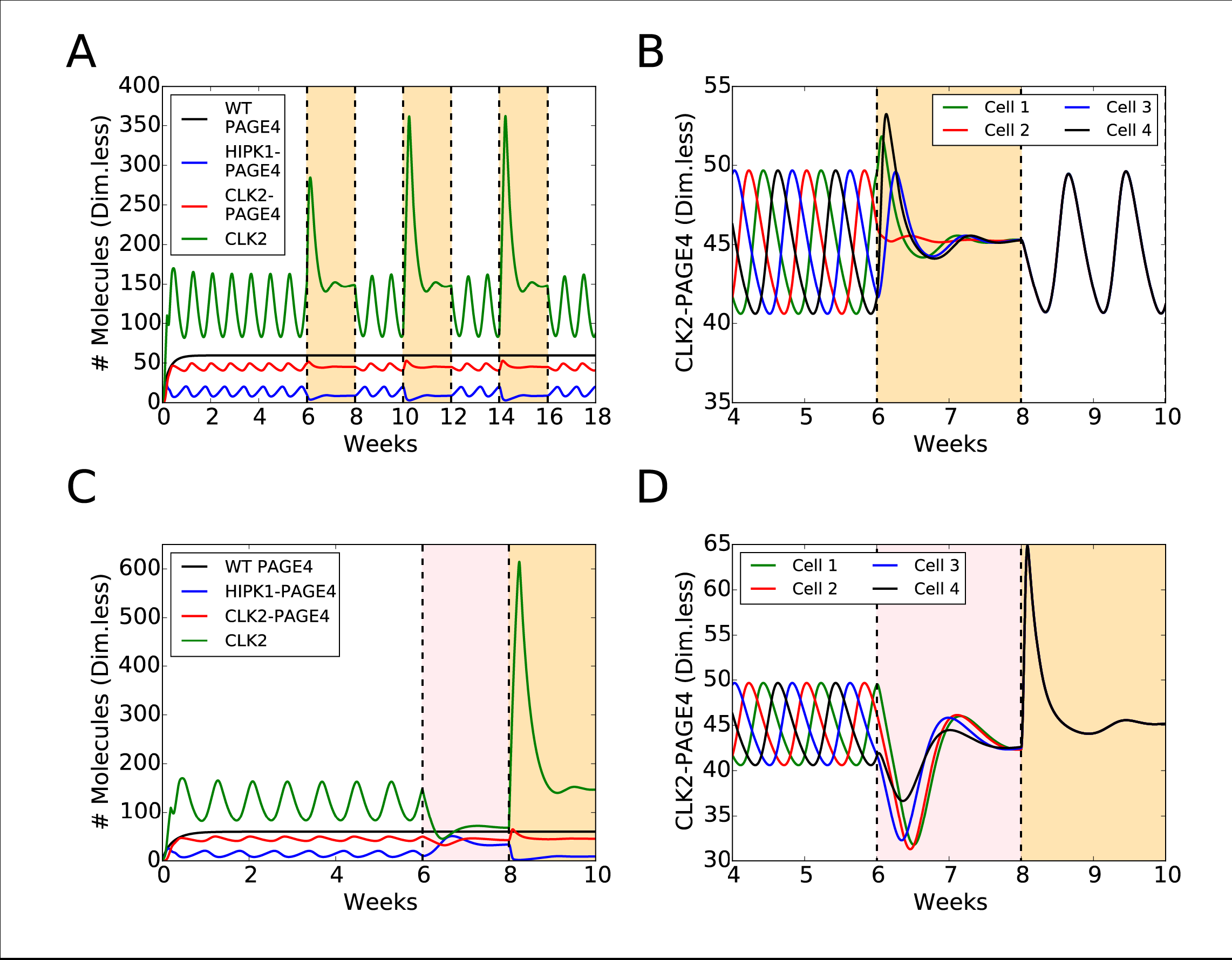
Intermittent ADT and BAT temporarily remove oscillations and synchronize the cells phenotypic dynamics. **A.** Temporal dynamics of WT PAGE4, HIPK1-PAGE4, CLK2-PAGE4 and CLK2 under Intermittent ADT. In the simulation, ADT was applied and removed at regular intervals of 2 weeks (the orange-shaded areas indicate when the therapy is on). **B.** Temporal dynamics of CLK2-PAGE4 in 4 different unsynchronized cells (i.e. the oscillations of CLK2-PAGE4 are not in phase). A 2-weeks pulse of ADT synchronizes the cells phenotypic dynamics. **C.** Temporal dynamics under BAT. The pink-shaded area indicates the 2 weeks of AR overexpression, while the orange-shaded area depicts the subsequent 2 weeks of ADT. Oscillations are quenched in 4 weeks of treatment, but the after 2 weeks, levels of HIPK1-PAGE4 decrease, while those of CLK2-PAGE4 increase. After 2 weeks of ADT, the levels are similar to those observed in continuous ADT. **D.** Similar to Intermittent ADT, BAT synchronizes the cell fates dynamics. In all plots, WT-PAGE4, HIPK1-PAGE4, CLK2-PAGE4 and CLK2 are represented in arbitrary dimensionless units.

Last, we considered the Bipolar Androgen Treatment (BAT), another novel treatment for prostate cancer [36]. This therapy considered 4 weeks of ADT, while an AR overexpression signal was on for the first 2 weeks. BAT suppressed oscillations during both treatment phases, but the cell was pushed to different states (Figure 6C). The activation of AR and the consequent inhibition of CLK2 enforced a higher level of HIPK1-PAGE4. Conversely, the last 2 weeks of ADT led to the already observed resistant phenotype with low HIPK1-PAGE4 (compare pink-shaded and orange-shaded regions in Figure 6C). Similar to the intermittent ADT case, BAT synchronized phenotypic oscillations of otherwise unsynchronized cells by quenching the oscillatory dynamics of PAGE4/AP-1/AR circuit (Figure 6D). Therefore, our results showed that similar to ADT, both BAT and IAD can also reduce phenotypic heterogeneity in a prostate cancer cell population, thus likely making the population more vulnerable to therapeutic strategies.

## 3 Discussion

It has become clear that a sizable fraction of eukaryotic proteins are unstructured before binding to a specific target [1]. The plasticity of these protein structures can facilitate the transcriptional and translational process, as well as the regulation of cellular functions. PAGE4 is a prototypical IDP with pleiotropic functions. Despite being considered almost entirely disordered based on data obtained using various biophysical measurements [38], in the present study, our simulations show that PAGE4 has functionally important structural features that are modulated by phosphorylation.

While the biophysical experimental techniques used previously to discern the various PAGE4 conformational ensembles are routinely used to study IDP, as far as we are aware, they have not been used in parallel to make measurements of the same IDP under identical experimental conditions. Furthermore, quite remarkably, the present data show a very high degree of similarity not only among the biophysical techniques used, but also between the theoretically computed and experimentally observed values.

Although the biophysical techniques employed can correctly capture some ensemble-based properties of PAGE4 such as the size and the distance in between two smFRET probes, there is still a gap between the highest time resolution that can be achieved experimentally and that of the structural dynamics of proteins. This problem is especially prominent for a protein that is highly disordered and exists as an ensemble of rapidly interconverting conformers. In this case, MD simulations with a physical force field can yield additional structural details and dynamical features of these proteins. In this study, our MD simulations accurately reproduce the ensemble averaged features of PAGE4 in both the unphosphorylated and phosphorylated states. This consistency provides good evidence that our model is able to correctly represent the structural ensemble seen experimentally.

Our studies also reveal the structural details underpinning the change of size of PAGE4 upon phosphorylation. Frequent N-terminal loop formation in both WT-PAGE4 and HIPK1-PAGE4, as reflected in our PCA analysis, is reminiscent of the “fly-casting” mechanism that has been proposed for other disordered proteins [31, 32, 33]. Here, a protein that typically has a relatively unstructured state can have a greater capture radius for a specific binding site than the folded state with its restricted conformational freedom. In this scenario of binding, it is envisioned that the unfolded state binds weakly at a relatively large distance followed by folding as the protein approaches the binding site, analogous to a fly-casting motion. The structural plasticity of the protein allows for a combination of binding and folding motions, which lowers the free-energy barrier for the protein-DNA recognition. In this study, PAGE4 is a highly charged molecule and therefore, it is possible that it can bind to both C-Jun of the AP-1 protein complex and DNA (the AP-1 binding site on the promoter). The regular dynamic motion of the N-terminal portion and the frequent loop formation can enlarge the scope for PAGE4 to find its binding partner, which would possibly increase the chance for the c-Jun transactivation. In contrast, CLK2-PAGE4 loses this regular motion with the replacement of a more disordered N-terminal motion. The loss of specificity may lead to a reduced binding affinity of PAGE4 [15] to AP-1, which eventually results in the degradation of PAGE4 by itself.

In addition, our simulations indicate an increase of turn-like structure in the central acidic region upon double- and hyperphosphorylation, and the loss of turn-like structure in the transient helical region upon hyperphosphorylation. Consistent with the former observation, a recent NMR study [26] also suggested that an increase of turn-like structure in the central acidic region was the source of a slightly more compact structure of HIPK1-PAGE4. The loss of turn structure in the transient helix indicates a partial order-disorder transition. The order-disorder transition has been previously shown to be a part of protein function to lower the free energy barrier and increase the kinetic rate of conformational rearrangements [39, 40, 41, 42]. Here, this transition offers another way by which phosphorylation modulates the binding affinity of PAGE4 to the AP-1 protein complex. Because the transient helical region was previously suspected to be a potential binding site of the AP-1 complex [26], this order-disorder transition can serve to disrupt the binding site, leading to the release of AP-1 upon PAGE4 hyperphosphorylation and the attenuation in c-Jun potentiation. Of note, this conjecture is based entirely on the structural analysis of PAGE4. Further studies including simulations for the equilibrated binding of PAGE4 to the AP-1 complex along with the cognate DNA binding site are currently underway.

To gain insight into the functional implications of the conformational dynamics of PAGE4 insofar as its role in state switching is concerned, we designed a coarse-grained mathematical model. The main goal of the model was to discern how the different conformational ensembles of PAGE4 may modulate androgen insensitivity in PCa cells through their opposite interactions with the AP- 1/AR axis. Our model predicted oscillations in the levels of the different PAGE4 conformational ensembles as well as HIPK1 and CLK2, the two kinases that are components of the PAGE/AP-1/AR regulatory circuit between an Androgen-Dependent (AD) and an Androgen-Independent (AI) phenotype. These oscillations correspond to a timescale of a week which corresponds approximately to the half-life of HIPK1-PAGE4. These observations support our previous experimental observations that cells in an isogenic population may exhibit phenotypes with varying androgen deprivation sensitivities [43].Therefore, it is plausible that, in addition to extrinsic signals, phenotypic plasticity can also arise from intracellular dynamics; in this case, by “conformational noise” [44] emerging from switching between different conformational states of PAGE4. Moreover, cells in a given population are highly likely to oscillate with some phase difference among each other, giving rise to non-genetic heterogeneity, a burgeoning therapeutic challenge in PCa [45, 46].

According to our model, all therapies — continuous Androgen Deprivation Therapy (ADT), Intermittent Androgen Deprivation (IAD) and Bipolar Androgen Therapy (BAT) — eliminated the oscillations and pushed cells toward a resistant phenotype. Furthermore, the treatments largely diminished phenotypic heterogeneity in a cohort of prostate cancer cells within 2 weeks. A limitation of our model is that we do not consider cell death being induced by any of the therapies; in other words, the model predictions mentioned above are only for a subset of cells that are not killed during therapeutic treatment. Extending this mathematical formalism to a population-level approach that considers accelerated cell death in the presence of hormonal treatments would represent a step forward toward formulating predictions of potential clinical relevance, especially given that ADT remains the standard of care for metastatic and locally advanced PCa for over 75 years. Oscillations were rescued when the inhibition on AR during IAD was removed. However, due to limited phenotypic heterogeneity, the cell population oscillated in phase after the treatment was applied. The perfect synchronization observed here can potentially also be disrupted by noise and biological effects beyond what is considered in the model. Yet, the hormonal treatment [47] could introduce a coherence time or window where the intracellular temporal dynamics of different cells appear correlated. Such predictions that can be experimentally validated using live-cell reporters for AR activity are currently being explored in our laboratories.

## 4 Materials and Methods

### 4.1 The AAWSEM Force Field

We used a recently developed model, Atomistic AWSEM (AAWSEM), for our simulation. AAWSEM is an adapted version of AWSEM. Details of AAWSEM have been elaborated elsewhere [23, 24]. Here we briefly summarize this force field. The AWSEM Hamiltonian is a coarse-grained force field that represents the backbone structure of the simulated proteins using three atoms per residue: *C_α_*, *C_β_* and O atoms. The positions of all the other backbone atoms can be calculated by assuming an ideal planar peptide bond geometry. The Hamiltonian can be decomposed into four components:

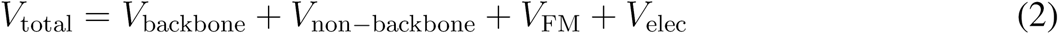

where *V*_non–backbone_ = *V*_contact_ + *V*_buriai_ + *V*_HB_. The *V*_backbone_ term dictates the backbone geometry of the protein. The *V*_non–backbone_ term describes the residue-residue interactions that are not related to backbone connectivity. The *V*_burial_ term is a many-body term that depends on the density of the immediate environs surrounding each residue. The *V*_HB_ hydrogen bonding term is governing the secondary structure of the protein. In practice, this term is correlated with the in-put based on secondary structure prediction [48, 25]. The *V*_contact_ = *V*_direct_ + *V*_Water_ is a contact term describing the tertiary interaction between two residues. It is composed of two terms. 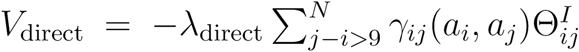 captures the direct protein-protein interaction. 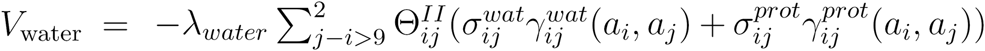 describes the water-mediated interaction. The *γ* parameters were optimized based on the training structures culled from the PDB database. The optimization was performed to maximize the ratio of the folding temperature to the glass transition temperature, *T_f_*/*T_g_* [49, 50, 51]. The fragment based associative memory term, V_FM_, is in the form of 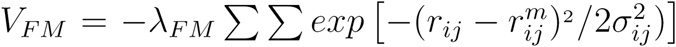, where σ_*ij*_ = |*i* — *j*|^0.15^ is a separation in protein sequence. *r_ij_* is a cartesian distance between the *C_α_* atoms of two residues i and j. 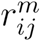 is the corresponding distance in the memory structures. V_FM_ guides the local-in-sequence interactions of the *C_α_* and *C_β_* atoms and can be based on fragment structures obtained in one of the two ways: In conventional AWSEM simulations, V_FM_ comes from existing PDB structures which share a similar local sequence with the simulated sequence. In AAWSEM, V_FM_ comes from exhaustive all-atom explicit-solvent simulations. Since disordered proteins do not usually have solved structures in the PDB database, here we choose to use atomistic simulations to obtain the fragment memory structures.

In the original version of AAWSEM, the full sequence of a protein was initially segmented in overlapping parts based on the output from secondary structure prediction tools such as PSSpred [25]. Next, the segments are simulated using an all-atom mode in explicit solvent. Here, since PAGE4 is predicted by PSSpred to be almost completely disordered, we segmented it according to the empirical motifs that have been annotated based on previous experiments [26]. The segmentation scheme is shown in Figure 1. Segments are simulated with an enhanced sampling method CST (Continuous Simulated Tempering) [27, 52] to achieve an equilibrated structural ensemble. The CST was performed in a temperature range of 293-350K and the structures that were sampled at temperatures below 330K were selected for a structural clustering. After the clustering of structures by the “single linkage” algorithm, the central structure was selected as the memory structure. The weight of the V_FM_ term is assigned according to the size of each structural cluster.

The *V*_elec_ electrostatic term is the Debye-Huckel term which approximates the screening effect of charge-charge interactions in solution. The details of this term were described in a separate paper [53]. Simply put, the form of *V*_elec_ is as follows:

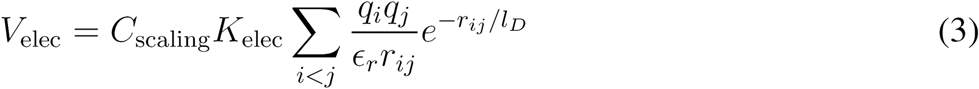

where *K*_Elec_ = (4πε_0_)^-1^ = 332.24 *kcal* · *Å* · *mol*^-1^ · *e*^-2^ and ε_r_ is the dielectric constant of the media. *q_i_* and *q_j_* are the charges of the residues. 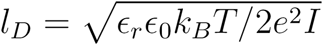 is the Debye length where *I* is the ionic strength of the solutions. Following the previous study, we use ε_r_ = 80.0 and *l_D_* = 10.0 Å throughout our simulations to achieve a typical physiological condition. *C*_scaling_ is a constant we introduced for tuning the relative strength of the electrostatic potential compared with other energy terms of the AWSEM Hamiltonian. In this study, we performed simulations with *C*_scaling_ = 1.0, 2.0 and 4.0. We found the simulations with *C*_scaling_ = 4.0 matched the experimental results quantitatively [15]. Smaller values of *C*_scaling_ resulted in qualitative agreement between simulations and experiments but poorer quantitative agreement (see the Supporting Information Figure S1, Table S1). Thus, the analyses throughout the main text are based on the *C*_scaling_ = 4.0 simulations. We used the same value of *C*_scaung_ to study different phosphorylated forms of PAGE4. Thus the changes in the conformations of PAGE4 upon phosphorylation are true predictions of this model. Also, we performed simulations in a neutral pH environment by setting the charges of Glu (Glutamic Acid) and Asp (Asparic Acid) residues to −1, while those of Arg (Arginine) and Lys (Lysine) residues were set to +1.

### 4.2 Treatment of Phosphorylation

Approaches to simulate phosphorylation have been studied in previous work using the AMH model [29, 30], which was the predecessor of AWSEM model. In that approach, the phosphorylated residues were simulated as “supercharged” Glutamic acid (Glu) residues. We adopted the same method in this study, where both the phosphorylated Ser (serine) and phosphorylated Thr (Threonine) were simulated as a Glus. Since the pKAs of phospho-serine and phospho-threonine were determined to be less than 7 in experiments [54], we simulated the phosphorylated residues with charge −2. Furthermore, because the phosphorylation probability was found to be variable among the different forms of PAGE4, we multiplied the charge by its phosphorylated probability to reflect this variability. For example, the Ser9 was shown by experiment to have a 30% ∼ 40% probability to be phosphorylated in HIPK1-PAGE4, so it was simulated as Glu with charge 35% × (−2) = −0.7. In the all-atom explicit-solvent simulations, the phosphoserine and phosphothreonine were simulated by adding phosphoryl groups onto Ser and Thr and simulated under CHARMM36 potential [55].

### 4.3 The Change of water mediated interaction for disordered proteins

The *γ* parameters in *V*_contact_ of the AWSEM potential were optimized in a self-consistent way to maximize the foldability of proteins [49, 50, 51]. In practice, the optimization is done by maximizing the *T_f_*/*T_g_*, or the ratio between the energy gap *δE* = *Aγ* and the roughness of the landscape Δ*E* = *γBγ*. Here A and *γ* are vectors and B is a matrix. The elements of A and B are defined as:

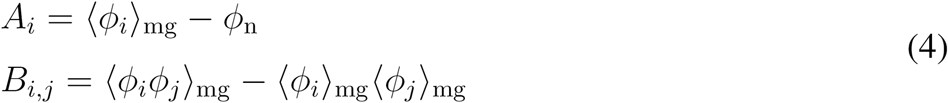

where *ϕ_i_* is the functional form of a particular type of interaction. The “mg” stands for “molten globule” and “n” stands for “native”. The optimization objective is the functional 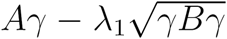, which is maximized with *γ* ∝ *B*^-1^*A*. In practice, *γ* parameters were determined in a self-consistent way by iterating through simulating to generate a set of molten-globule structural decoys and recalculating its value until *γ* converges. Additionally, The collapse temperature of the simulated proteins is related to *γ* parameters by *T*_c_ = *A’γ*, whose element is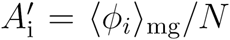 and *N* is the number of residues. The degree of collapse of proteins in a simulation with a fixed temperature can be controlled by changing *T_c_*. Therefore, we can change the degree of collapse of simulated proteins by shifting the *γ* parameters.

Our simulation with the default model showed an over-compact structure of PAGE4 (Table S2 and Figure S2). To resolve this problem, we shifted the *γ* parameters of *V*_contact_ to make the AWSEM potential, on average, more repulsive. The shifting number used throughout our simulations was –0.7, which ensured the simulated ensemble of WT-PAGE4 has a similar *R_g_* to that measured from the SAXS experiments [15]. We used the same *γ* values in all of our simulations. In this way, any difference of *R_gs_* among three phosphoforms of PAGE4 from our simulations comes from the effect of phosphorylation.

### 4.4 Details of Atomistic Simulations

The atomistic simulations were performed for the segments of PAGE4 in an explicit-solvent environment (Figure 1). We use the CST [27] method to speed up the thermodynamic sampling. The temperature was continuously changing ranging from *T* = 293 ∼ 350K, and the detailed balance was kept throughout the simulations. The atomistic simulation were carried out with a time step of 2.0 fs. The simulated system was neutralized in a physiological condition (ionic concentration of 0.15M) by Na_+_ and Cl^−^ ions. For each segment, we performed a total of 1 μs of atomistic simulation with CHARMM36m force field, which was calibrated based on the latest forcefield of CHARMM36 for IDPs [56]. After the simulations, we used only data with *T* < 330K to feed into the AWSEM V_FM_ potential. We also disregarded the first 25 ns of each simulation, which was required for convergence of the CST algorithm.

### 4.5 Mathematical Model for PAGE4/AP-1/AR regulartory axis

We modelled the PAGE4 circuit depicted in Figure 5A by extending the mathematical framework formulated by Kulkarni et al [15]. The temporal dynamics of WT-PAGE4 (*P_U_*), HIPK1-PAGE4 (*P_M_*), CLK2-PAGE4 (*P_H_*) and the CLK2 enzyme (*C*) in the prostate cancer cell is described by:

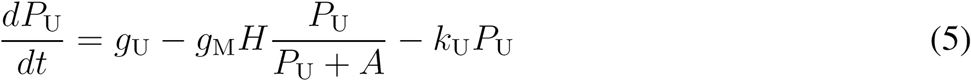

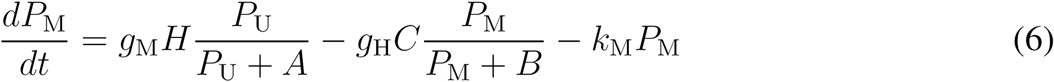

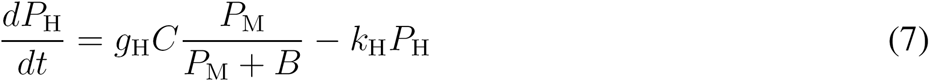

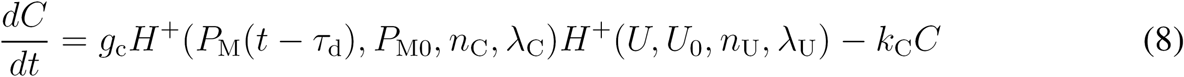

where (*g_U_,g_M_,g_H_,g_C_*) and *k_U_, k_M_, k_H_, k_C_*) are basal production and degradation rates. The term 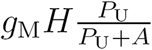 represents the conversion of WT-PAGE4 (*P_U_*) to HIPK1-PAGE4 (*P_M_*) via enzymatic activity of Hipk1 (*H*). Here, A represents the threshold level for the catalytic reaction. Similarly 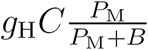 represents the conversion of HIPK1-PAGE4 (*P_M_*) to CLK2-PAGE4 (*P_H_*) via the enzyme CLK2 (C) and threshold *B*. *P_M_* regulates *C* via the intermediates C-Jun and AR activity. This interaction is modelled via the shifted Hill function [57] 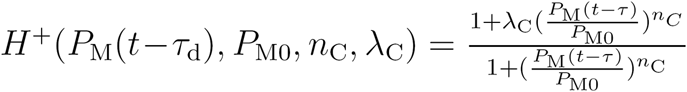 where λ_C_ is the production rate fold-change of *C* in the presence of HIPK1-PAGE4, *n_C_* is the Hill coefficient regulating the fold-change dependence on *P_M_*, *P_M0_* is a threshold concentration for *C* activation and *P_M_* (*t* — τ_d_) is the level of *P_M_* at time *t* — τ_d_. The delay term *τ_d_* considers the intermediate steps in the signaling.

The androgen treatments effectively regulate CLK2 via the modulation on AR activity (see Figure 5A), and were therefore modelled with the term *H^+^(U,U_0_,*n*_U_*, λ_u_). *U*, *U*_0_, *n_u_*, λ_u_ are the treatment level, the threshold for treatment activation, the Hill coefficient and the fold-change in production rate of CLK2. The fold-change for CLK2 production rate was λ_u_ > 1 for ADT (inhibition of AR and, therefore, activation of CLK2) and λ_u_ < 1 for the overexpression phase of BAT (overexpression of AR, inhibition of CLK2).

A dimensionless version of Equation (5-8) was used for the numerical solution of the model. The Supplementary Information provides details about the dimensionless model (Section “Dimensionless model”), the model’s parameters (Section “Parameters estimation”) and the numerical solution (Section “Details on numerical simulation”).

## 5 Acknowledgement

This work is supported by the Center for Theoretical Biological Physics sponsored by the NSF (Grant PHY-1427654) and by NSF-CHE 1614101. XL was partially supported by the National Institute of Health Grant R01-GM110310. We also wish to acknowledge support from NIH grants GM062154 (JO) and CA181730 (JO and PK). GR was supported by JC Bose Fellowship and SERB Centre for Mathematical Biology Phase II grant. MKJ was supported by a Gulf Coast Consortia on the Computational Cancer Biology Training Program (CPRIT Grant No. RP170593).

